# duphold: scalalable, depth-based annotation and curation of high-confidence structural variant calls

**DOI:** 10.1101/465385

**Authors:** Brent S. Pedersen, Aaron R. Quinlan

## Abstract

Most structural variant detection tools use clusters of discordant read-pair and split-read alignments to identify variants, yet do not integrate depth of sequence coverage as an additional means to support or refute putative events. Here, we present *duphold*, as a new method to efficiently annotate structural variant calls with sequence depth information that can add (or remove) confidence to SV predicted to affect copy number. It indicates not only the change in depth across the event, but also the presence of a rapid change in depth relative to the regions surrounding the breakpoints. It uses a unique algorithm that allows the run time to be nearly independent of the number of variants. This performance is important for large, jointly-called projects with many samples, each of which must be evaluated at thousands of sites. We show that filtering on *duphold* annotations can greatly improve the specificity of deletion calls and that its annotations match visual inspection. Duphold can annotate structural variant predictions made from both short-read and long-read data. It is available under the MIT license at: https://github.com/brentp/duphold.

## Findings

### Motivation

Structural variants (SV) are a broad class of genetic variation including duplications, deletions, inversions, insertions, and translocations. SVs are known to be more difficult to detect with high accuracy than single-nucleotide and insertion-deletion variants. As such, the false positive rate can be high, so it is difficult to separate true variants of interest from those that might be due to noise. The most commonly used structural variant callers^1–5^ use two types of sequence alignments to discover structural variation: paired-end reads having an unusual orientation or insert size (so called “discordant pairs”), and split-reads, where the sequence is aligned to different parts of the genome. These methods work well, but do not use the aligned sequence depth to detect or filter structural variant calls. This is an important limitation, since, for example, we expect a true hemizygous deletion to exhibit 50% of the sequence coverage of flanking diploid regions. Our experience in evaluating thousands of candidate SVs with SVPlaudit^6^, which allows users to quickly assess the veracity of structural variants, we noted two consistent patterns that distinguished confident deletion and duplication calls from apparent false positives. First, events without an obvious reduction or increase in coverage are much less likely to appear as “real” events to the human eye. Second, events with a rapid change in depth at (or near) the breakpoints are more plausible. Obvious false positive calls lack either or both of those signals. Unfortunately, most SV discovery tools do not look at the coverage information required to discover these signals. We therefore developed *duphold* to enforce the observations we made through manual inspection and rapidly annotate SV calls in order to prioritize high-quality variant calls.

### Implementation

*duphold* uses our previously described hts-nim^7^ library to quickly extract coverage information from a BAM or CRAM file into an array where it can be queried directly. These depth profiles are used to quickly annotate a VCF^8^ file of structural variants with coverage calculated from a BAM or CRAM file of alignments. Briefly, *duphold* allocates an (int16) array whose size is the length of the current chromosome (this array uses about 500 megabytes of memory for the 249 megabase human chromosome 1), iterates over each read in a BAM or CRAM for that chromosome, and increments any bases where a read (or segment of a read) starts and decrements any bases where a read (or part of a read) ends. A segment of a read is defined by the SAM^9^ CIGAR operations. Once *duphold* has processed all segments for all alignments in a chromosome, it performs a cumulative sum which results in a per-base coverage value in the array. A 64 bit integer is used to track the actual depth but the depth stored on the arrays is capped at at the maximum value for a 16 bit integer (32767) to prevent integer overflow. This algorithm is fully detailed in the mosdepth manuscript^10^. Once the coverage array is filled, all remaining steps are independent of the number of alignments. Owing to the speed of in-memory array operations, subsequent depth calculations are nearly independent of the number of variants annotated in the VCF file.

For each structural variant, *duphold* annotates the VCF sample format field of the variant with both the change in depth relative to the surrounding 5000 bases on either side of the event, and the fold-change in coverage in the breakpoint relative to other regions in the genome with similar GC-content. In order to compare the coverage observed for each variant with genomic bins of similar GC-content, duphold calculates the GC-content in each 250 base window (the window size is configurable) in the chromosome along with the median depth in that window. This requires 0.55 CPU-seconds for chromosome 1. These per-window depth and GC values are used as a distribution against which to compare incoming variants.

Once the depths and the GC-windows are calculated, we use hts-nim to read and annotate structural variant calls in VCF format. For each variant, the GC-content from start to end is calculated, and the median depth inside the event is compared to the window values with a similar GC-content to calculate a fold-change value (DHBFC for duphold bin fold-change). *Duphold* then compares the median depth in the event to the median depth from the 5000 bases on either side; this measure (named DHFFC, for duphold flank fold-change) captures the change in depth one would observe by eye upon visual inspection. The depth fold-change values are added to the sample’s format information in the variant’s VCF entry. *Duphold* is run on a single-sample at a time, but it has support for facile parallelization across samples. It can run on a 25X whole genome CRAM in < 15 CPU-minutes.

### Evaluation

#### Deletions

We evaluated *duphold* by annotating the lumpy^1^ calls and svtyper^11^ genotypes we produced for the HG002 sample sequenced by the Genome in a Bottle^12^ (GiaB). We compared these to the GiaB truth-set of deletions for the same sample. We used the *duphold* annotations to filter to more stringent call sets and evaluate the precision and recall. Because *duphold* does not add any new variants, it can only improve precision, not recall.

The duphold depth annotations enable simple filters that reduce the number of false-positives while retaining most true positives (Table 1). For example, requiring that the fold-change of the deletion relative to the 5000 bases flanking the deletion must be less than 0.7 (DHFFC < 0.7) removes only 85 of the original 133 false positive calls (64%), while retaining 1817 of the original 1849 true positive calls (98%). The DHBFC metric measures the depth fold-change relative to bins with a similar GC-content, and performs similarly. Using more stringent filtering can further reduce the false positive rate at the expense of the recall. The information used in this filtering is independent of the values reported by lumpy and svtyper which do not look at sequence depth metrics.

**Table 1.** Evaluating accuracy of calls filtered by duphold annotations. We evaluated deletion calls from lumpy+svtyper using truvari.py (https://github.com/spiralgenetics/truvari) with the GiaB v0.6 truthset. Columns are FDR: false discovery rate, FN: false negatives, FP: false positive, TP: true-positive, precision, recall, and F1 score. This shows that using either the DHBFC < 0.7 or DHFFC < 0.7 as a filtering criteria for deletions increases precision, removing 64% (1 - 48 / 133)) of false positive calls while retaining about 98% (1817 / 1849) of true positive calls in the case of using DHFFC.

| Method | FDR | FN | FP | TP-call | Precision | Recall | F1-score |
| --- | --- | --- | --- | --- | --- | --- | --- |
| Unfiltered | 0.067 | 900 | 133 | 1849 | 0.933 | 0.673 | 0.782 |
| DHBFC < 0.7 | 0.023 | 944 | 43 | 1805 | 0.977 | 0.657 | 0.785 |
| DHFFC < 0.7 | 0.026 | 932 | 48 | 1817 | 0.974 | 0.661 | 0.788 |

We examined each of the 48 false positive calls that remained after *duphold* filtering. These included a mixture of complex regions that had a loss of coverage, and some that looked like they could be real variants, but with minimal alignment support. We also visually inspected each of the 32 (i.e., 1849 - 1817) true positives that *duphold* marked as low confidence owing to a flank fold-change greater than 0.7 (DHFFC > 0.7). Most of these had a minimal change in coverage that did not meet our threshold and many looked like they did not have strong evidence for a call. We even noted one variant that looked like a duplication within a deletion, resulting in a copy-neutral event. While these highlight the limitations of a purely depth-based approach, we find that the more than 2-fold reduction in false positives in concert with a retention of nearly 99% of true-positives to be a convincing demonstration of *dupholds*’s power to remove the abundant false positive SV prediction common to most analyses.

### Scaling

We designed *duphold* with the expectation that it would be used on large datasets where specificity and run time are both critical. For this reason, we optimized it for situations where it would be used to evaluate many thousands of variants. In an effort to measure scaling performance, compared the times of both svtyper and duphold on subsets of the thousand genomes phase 3 structural variants (Figure 1). We are not interested in the direct time comparison with svtyper (since svtyper does more work to genotype the variants). Instead, the relevant pattern is the trajectory in order to demonstrate how well *duphold* scales. Svtyper follows a linear increase in time with the number of variants, while *dupholds’s* performance is nearly independent of the number of variants, using either a single (green) or 3 (red) threads. This performance is driven by the fact that all of the alignment data is read into efficient data structures that can be queried thousands of times a second. This strategy incurs a large initial cost to construct the data structure, and therefore makes *duphold* less efficient for small variant sets. We have chosen to optimize for larger variant sets, since this context is where efficiency is most important.

**Figure 1.**
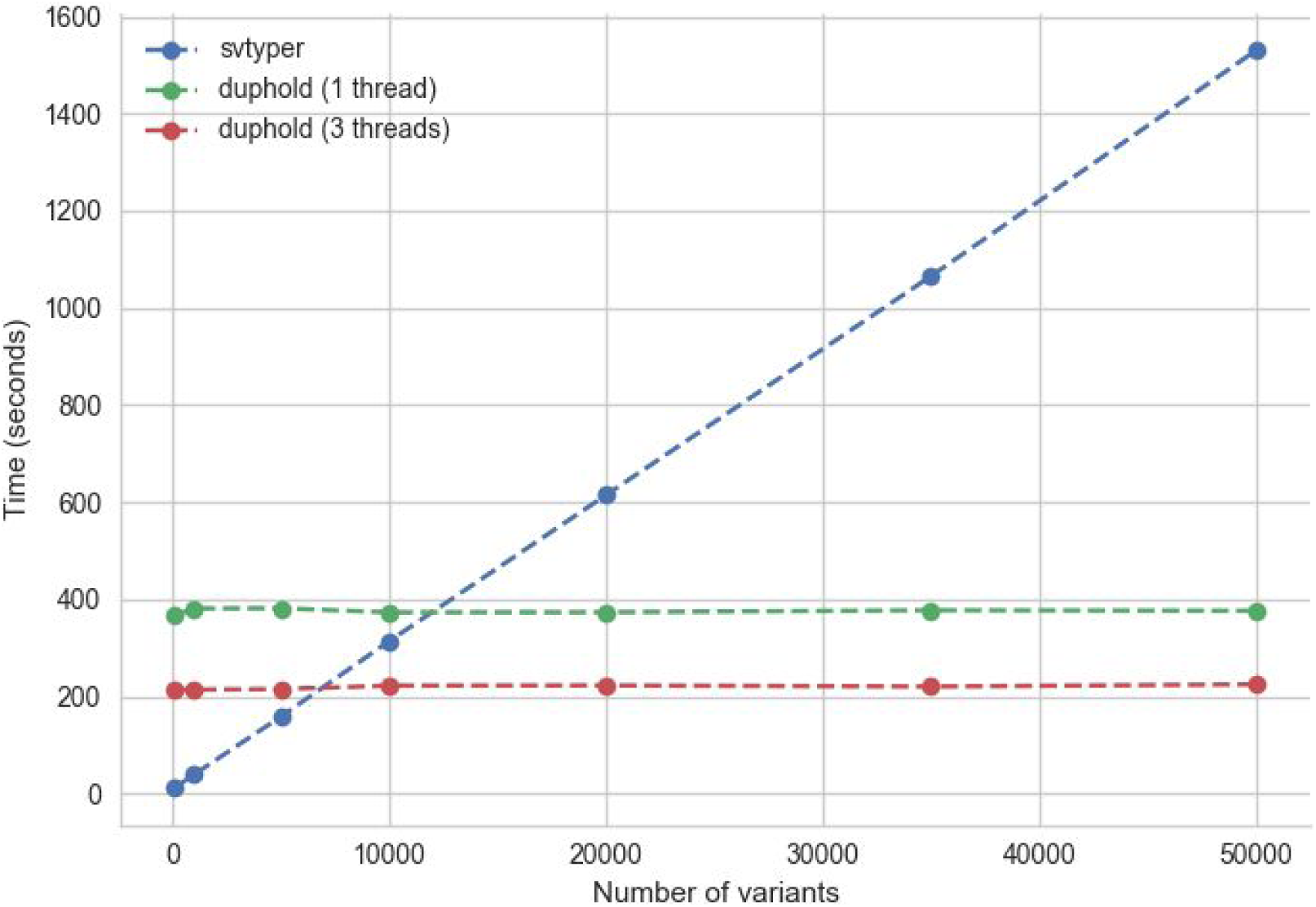
Duphold scalability. The time to annotate (or genotype) for duphold and svtyper is shown (y-axis) as a function of the number of variants tested (x-axis). While svtyper (blue) exhibits a linear increase in type with the number of variants, *duphold* is relatively independent of the number of variants. There is an initial cost that makes the *duphold* strategy less efficient for few (less than about 10K) variants but it scales well to annotating thousands of variants as we expect for large cohorts.

## Methods

To evaluate the ability of *duphold* to prioritize structural variant calls, we used data from the Genome in a Bottle project for sample HG002. We downloaded all fastqs from: ftp://ftp-trace.ncbi.nlm.nih.gov/giab/ftp/data/AshkenazimTrio/HG002_NA24385_son/NIST_HiSeq_HG002_Homogeneitv-10953946/HG002_HiSeq300x_fasta/140528_D00360_0018_AH_8VC6ADXX/,aligned with bwa-mem^13^, and marked duplicates with samblaster^14^ to generate a CRAM file with ~25X median sequence coverage. We used the GiaB SV calls from ftp://ftptrace.ncbi.nlm.nih.gov/giab/ftp/data/AshkenazimTrio/analvsis/NIST_SVs_Integration_v0.6/ as our truth-set. Because that does not explicitly differentiate insertions from duplications. we limited our evaluation to duplications. We ran lumpy^1^ and svtyper^11^ via smoove (https://github.com/brentp/smoove) to create and genotype structural variant calls. We evaluated the precision and recall before and after applying various filtering on the duphold annotated variants using truvari (https://github.com/spiralgenetics/truvari). We used samplot (https://github.com/ryanlayer/samplot) to look at individual variants that were called as true positives, false positives and false negatives.

The truvari command used was:

~~~
truvari.py -s 300 -S 270 -b HG002_SVs_Tier1_v0.6.DEL.vcf.gz -c $lumpy_vcf
-o eval-no-support --passonly --pctsim=0 -r 20 --giabreport -f $fasta
--no-ref --includebed HG002_SVs_Tier1_v0.6.bed -O 0.6
~~~

To demonstrate the utility of *duphold* on duplication calls, we downloaded the sniffles^15^ SV calls from http://labshare.cshl.edu/shares/schatzlab/www-data/fsedlaze/Sniffles/GiaB/all_reads.fa.giab_h002_ngmlr-0.2.3_mapped.bam.sniffles1kb_auto_noalts.vcf.gz and annotated DUP calls in that VCF with *duphold* using the same Illumina HG002 cram file as above.

To evaluate the scaling on realistic sites, we used duphold to annotate the same HG002 file, but on the 68,818 variants from the 1000 Genomes SV calls at: ftp://ftp.1000genomes.ebi.ac.uk/vol1/ftp/phase3/integrated_sv_map/ALL.wgs.mergedSV.v8._20130502.svs.genotypes.vcf.gz. We limited those calls to the variants that could be genotyped by svtyper (excluding insertions). We then randomly chose 100, 1000, 10K, 20K, 35K and 50K variants and ran svtyper and duphold on each set. We also ran duphold with 3 threads to evaluate the benefit of parallelization.

We downloaded the HG002 SNP/Indel calls from: ftp://ftp-trace.ncbi.nlm.nih.gov/giab/ftp/release/AshkenazimTrio/HG002_NA24385_son/latest/GRCh37/

## Conclusions

*Duphold* enables rapid annotation of existing structural variant calls with sequence depth information that facilitates the distinction between high and low confidence deletions and duplications. Using the Genome in a Bottle truth set, we have shown that we can exclude nearly 64% of false positives SV predictions while retaining over 98% of true positive variants using a simple filter on a *duphold* annotated VCF. Given the minimal additional runtime of as few as 25 minutes for a 30X genome, this is a substantial improvement for the overall accuracy of SV callsets.

## Availability of supporting source code and requirements

Project name: duphold

Project home page: https://github.com/brentp/duphold

Operating system(s): binary available for linux (can be built on OSX and windows) Programming language: nim

Other requirements: htslib.so >= 1.8

License: MIT

## Declarations

## List of abbreviations

GiaB: genome in a bottle
SNP: single nucleotide polymorphism
SV: structural variant
VCF: variant call format

## Consent for publication

Not applicable

## Author’s contributions

BSP designed and wrote the software, performed the analyses and co-wrote the paper. ARQ co-wrote the paper.

